# BLOCKADE OF M4 MUSCARINIC RECEPTORS ON STRIATAL CHOLINERGIC INTERNEURONS NORMALIZES STRIATAL DOPAMINE RELEASE IN A MOUSE MODEL OF *TOR1A* DYSTONIA

**DOI:** 10.1101/2020.12.17.423273

**Authors:** Anthony M. Downs, Yuping Donsante, H.A. Jinnah, Ellen J. Hess

## Abstract

Trihexyphenidyl (THP), a non-selective muscarinic receptor (mAChR) antagonist, is commonly used for the treatment of dystonia associated with *TOR1A*, otherwise known as *DYT1* dystonia. A better understanding of the mechanism of action of THP is a critical step in the development of better therapeutics with fewer side effects. We previously found that THP normalizes the deficit in striatal dopamine (DA) release in a mouse model of *TOR1A* dystonia (*Tor1a*^*+/ΔE*^ knockin (KI) mice), revealing a plausible mechanism of action for this compound, considering that abnormal DA neurotransmission is consistently associated with many forms of dystonia. However, the mAChR subtype(s) that mediate the rescue of striatal dopamine release remain unclear. In this study we used a combination of pharmacological challenges and cell-type specific mAChR conditional knockout mice of either sex to determine which mAChR subtypes mediate the DA release-enhancing effects of THP. We determined that THP acts in part at M4 mAChR on striatal cholinergic interneurons to rescue DA release in both *Tor1a*^*+/+*^ and *Tor1a*^*+/ΔE*^ KI mice. Further, we found that the subtype selective M4 antagonist VU6021625 recapitulates the effects of THP. These data implicate a principal role for M4 mAChR located on striatal cholinergic interneurons in the mechanism of action of THP and suggest that subtype selective M4 mAChR antagonists may be effective therapeutics with fewer side effects than THP for the treatment of *TOR1A* dystonia.

## INTRODUCTION

Dystonia associated with *TOR1A* variants, otherwise known as *DYT1* dystonia, is an inherited form of dystonia caused by a three base pair deletion (Δgag) in the *TOR1A* gene (Ozelius et al., 1997). Because the precise mechanism(s) that lead to the abnormal movements are unknown, treatments for *TOR1A* dystonia are inadequate. The non-selective muscarinic acetylcholine receptor (mAChR) antagonist trihexyphenidyl (THP) is the preferred oral pharmaceutical for *TOR1A* dystonia, but its clinical use is limited due to significant dose-limiting side effects associated with its nonselective action at all five mAChRs subtypes (Schwarz and Bressman, 2009; Thenganatt and Jankovic, 2014). A better understanding of the mAChR subtype(s) responsible for the therapeutic effect of THP is critical for the development of targeted treatments with fewer side effects.

One potential mechanism of action of THP is the regulation of striatal dopamine (DA) neurotransmission. In patients, both DA metabolites and D2 DA receptor availability are abnormal (Augood et al., 2004; Asanuma et al., 2005). In mouse models, striatal DA release is reduced (Balcioglu et al., 2007; Page et al., 2010; Song et al., 2012; Downs et al., 2019). Striatal DA release is regulated by acetylcholine (ACh) (Zhang et al., 2002b; Zhang et al., 2002a; Exley and Cragg, 2008; Zhang et al., 2009a; Zhang et al., 2009b; Threlfell et al., 2010) and we recently discovered that THP rescues the deficit in both evoked DA release and steady-state extracellular DA concentrations in the striatum of a mouse knockin (KI) model of *TOR1A* dystonia (*Tor1a*^*+/ΔE*^ KI mice) (Downs et al., 2019). However, the mAChR subtype(s) and striatal cell type that mediate the effects of THP are unknown.

THP binds to all 5 mAChRs with nanomolar affinity (Dorje et al., 1991; Bolden et al., 1992), and all 5 mAChRs are expressed in the striatum (Buckley et al., 1988; Levey et al., 1991; Harrison et al., 1996; Alcantara et al., 2001; Yamada et al., 2003; Hernandez-Flores et al., 2015). M1 mAChRs are expressed on both direct and indirect striatal projection neurons (dSPNs and iSPNs) where they mediate corticostriatal plasticity (Harrison et al., 1996; Hernandez-Flores et al., 2015). M1 mAChRs also regulate extracellular DA concentrations, although the underlying mechanisms are unknown (Gerber et al., 2001a). M4 mAChRs are expressed on dSPNs where they regulate corticostriatal plasticity (Nair et al., 2015; Nair et al., 2019) and mediate striatal DA release via endocannabinoid signaling (Foster et al., 2016). Additionally, M4 mAChRs expressed on cholinergic interneurons (ChIs) act as inhibitory autoreceptors to modulate extracellular ACh concentrations, which play a central role in mediating DA release (Threlfell et al., 2010; Pancani et al., 2014). While M2 and M3 mAChRs are located on glutamatergic axon terminals within the striatum (Buckley et al., 1988; Levey et al., 1991; Alcantara et al., 2001), we have previously demonstrated that THP does not depend on glutamatergic signaling to modulate striatal DA release (Downs et al., 2019). M5 mAChR are expressed exclusively on DA terminals in the striatum (Yamada et al., 2003). However, previous studies have demonstrated that activation of M5 mAChRs in the striatum stimulates DA release, so it is unlikely that the mAChR antagonist THP enhances DA release through this mechanism (Foster et al., 2014). Therefore, THP likely acts through M1 or M4 mAChRs or both to enhance DA release. In this study, we used a combination of pharmacologic challenges and cell-type specific mAChR knockout (KO) mice to determine the role of M1 and M4 mAChRs in the potentiation of DA release by THP in a KI mouse model of *TOR1A* dystonia (*Tor1a*^*+/ΔE*^).

## MATERIALS AND METHODS

### Animals

Male and female mice (8-14 weeks of age) inbred on C57BL/6J were used for all studies. All mice were bred at Emory University. Animals were maintained on a 12h light/dark cycle and allowed *ad libitum* access to food and water. All experimental procedures were approved by the Emory Animal Care and Use Committee and followed guidelines set forth in the *Guide for the Care and Use of Laboratory Animals*. Heterozygous knockin mice carrying the *Tor1a(Δgag)* mutation (*Tor1a*^*+/ΔE*^) (Goodchild et al., 2005) and normal littermates (*Tor1a*^*+/+*^) were genotyped using PCR (forward primer 5’-GCTATGGAAGCTCTAGTTGG-3’; reverse primer 5’- CAGCCAGGGCTAAACAGAG-3’). The following strains were used for the conditional mAChR KO mice experiments (Table 1): M4 mAChR flox (M4^flox/flox^) (Jeon et al., 2010), M1 mAChR flox (M1^flox/flox^) (Kamsler et al., 2010), *ChAT-cre* (ChAT^tm1(cre)/Lowl^/MwarJ; JAX #031661), *D1-cre* (Tg(Drd1-cre)^EY262Gsat^/Mmucd; MMRRC #030989-UCD), and *A2A-cre* (Tg(Adora2a-cre)^KG139Gsat^/Mmucd; MMRRC #036158-UCD). These strains were used to generate both *Tor1a*^*+/+*^ and *Tor1a*^*+/ΔE*^ mice with conditional KO of M4 mAChR from cholinergic neurons (ChAT-cre; M4^flox/flox^), KO of M4 mAChR from dSPNs (D1-cre; M4^flox/flox^), KO of M1 mAChR from dSPNs (D1-cre; M1^flox/flox^), and KO of M1 from iSPNs (A2A-cre; M1^flox/flox^). Mice were genotyped using PCR with the following primers *D1-cre* (forward primer 5’- GCTATGGAGATGCTCCTGATGGAA-3’; reverse primer 5’- CGGCAAACGGACAGAAGCATT -3’); *A2A-cre* (forward primer 5’- CGTGAGAAAGCCTTTGGGAAGCT-3’; reverse primer 5’- CGGCAAACGGACAGAAGCATT-3’); *ChAT-cre* (WT forward primer, 5’-GCAAAGAGACCTCATCTGTGGA-3’; cre forward primer, 5’- TTCACTGCATTCTAGTTGTGGT-3’; common reverse primer, 5’- GATAGGGGAGCAGCACACAG-3’); *M1 flox* (WT forward primer 5’- GAGCCTCAGTTTTCTCATTGG-3’; mutant forward primer 5’- AACACTACTTACACGTGGTGC- 3’; common reverse primer 5’- TCAACCTGTACTGGTGATACG-3’); *M4 flox* (forward primer 5’- TGCAGATGTAGCTCAGCTCAGCGGTAC-3’; reverse primer 5’- TGAAGGTTGTAGACAAAGCTATACACATGGC-3’). Controls for the conditional KO experiments were homozygous for the flox’d muscarinic receptor allele but did not carry a cre transgene. All strains were maintained on a C57BL/6J background.

**Table 1.**
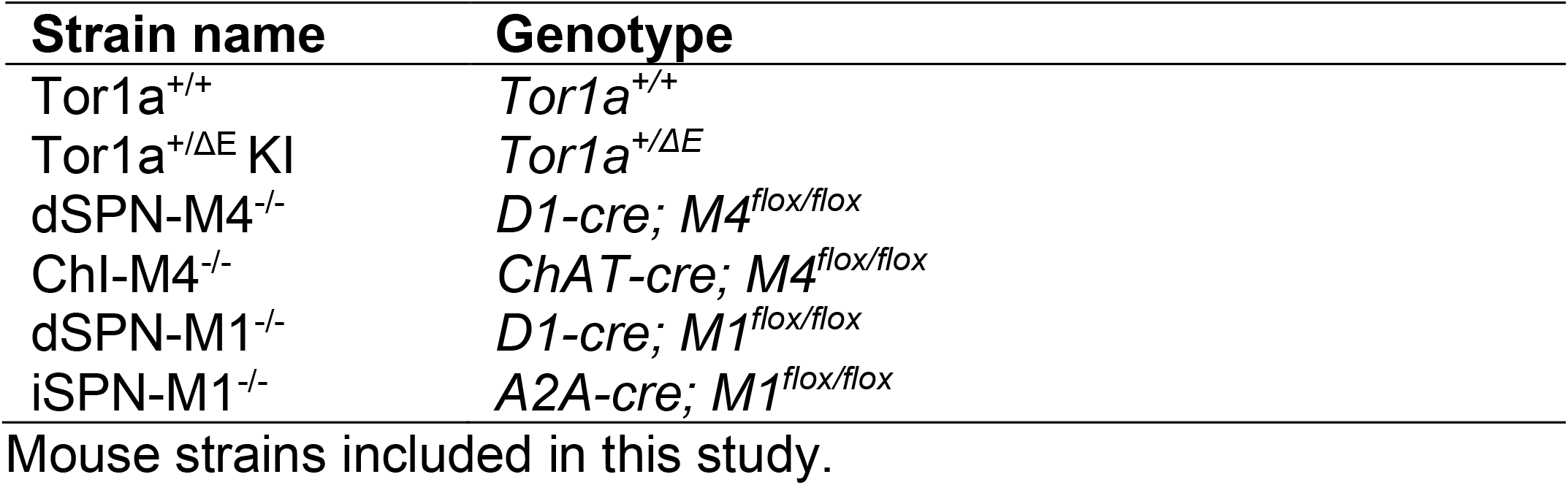
Mouse strains and genotypes

### Fluorescent in situ hybridization

Brains were rapidly dissected, frozen and kept at -80°C until sectioning (10 μm). RNAscope fluorescent *in situ* hybridization was performed according to manufacturer’s instructions (Advanced Cell Diagnostics, Newark, CA, USA). Briefly, slides were fixed in ice-cold 4% paraformaldehyde in 0.1M PBS (phosphate buffered saline) for 15 mins. Slides were then dehydrated in graded ethanols and allowed to air dry for 5 minutes. Slides were treated with Protease-IV solution for 30 mins at room temperature and then incubated with the appropriate probes for 2 hours at 40°C. The following probes were used: Mm-*Chat*-C1 (cat # 408731), Mm-*Chrm4*-C2 (cat # 410581-C2), Mm-*Drd1*-C3 (cat # 461901-C3), Mm-*Chrm1*-C1 (cat # 495291), Mm-*Adora2a*-C2 (cat # 409431-C2). A series of four amplification steps were then performed according to manufacturer’s instructions with 2 × 2 min washes in wash buffer between each amplification step. Slides were counterstained with DAPI and then coverslipped using Prolong Glass Antifade Mounting media (Thermo Fisher Scientific, Waltham, MA, USA). Slides were stored at 4°C until imaging.

Slides were imaged using a Leica SP8 confocal microscope using Leica Acquisition Suite (LAX S) (Leica Microsystems, Buffalo Grove, IL, USA). A total of 6 regions of interest (ROIs) from the dorsolateral quadrant of the striatum were taken from each section using a 40X or 60X objective and an optical thickness of ∼5 μm. The same optical settings were used for image acquisition from each mouse with the settings titrated for each specific probe. Quantification of labeled cells was performed using the Cell Counter function in ImageJ (NIH). For each genotype, only cells that were labeled with RNAscope probes were counted, rather than counting all DAPI-positive cells in the striatum. Data were presented as both percentage of positively labeled cells for each marker as well as the number of cells for each probe. Cells were considered positive for a given probe if there were >5 particles clustered around the nucleus.

### Fast scan cyclic voltammetry

Fast scan cyclic voltammetry (FSCV) to measure DA release was performed according to previously published methods (Downs et al., 2019). Mice were euthanized using cervical dislocation and 300 μm sections were prepared using a vibratome in ice-cold oxygenated sucrose-supplemented artificial cerebral spinal fluid (aCSF) containing [in mM]: sucrose [194], NaCl [20], KCl [4.4], CaCl_2_ [1.2], MgCl_2_ [1.2], NaH_2_PO_4_ [1.2], NaHCO_3_ [25], D-glucose [11] at pH 7.4. After sectioning, brain slices were maintained in a holding chamber containing oxygenated bicarbonate-buffered aCSF containing [in mM]: NaCl [126], KCl [2.45], CaCl_2_ [2.4], MgCl_2_ [1.2], NaH_2_PO_4_ [1.2], NaHCO_3_ [25], D-glucose [11] and maintained at room temperature for 45-60 min before experiments began.

All FSCV experiments were performed in the dorsolateral striatum at ∼ Bregma +0.26 mm because this region receives dense innervation from the motor cortex (Hintiryan et al., 2016). A slice was transferred to the recording chamber and perfused with oxygenated aCSF at 32°C for 30 min to equilibrate to the chamber. A carbon fiber electrode constructed in-house was inserted approximately 50 μm into the surface of the slice and a bipolar tungsten stimulating electrode was placed approximately 200 μm away. DA release was evoked by a 1-pulse (600 μA, 4 ms pulse width) electrical stimulation at 5 min inter-stimulus intervals to prevent rundown. The scan rate for voltammetry was 400 V/s from -0.4 V to 1.3 V to -0.4 V verses Ag/AgCl with a sampling rate of 10 Hz using a Chem-Clamp voltammeter-amperometer (Dagan Corporation, Minneapolis, MN, USA). FSCV experiments were conducted and analyzed using Demon voltammetry software (Wake Forest University) (Yorgason et al., 2011) to determine peak DA release. All compounds were diluted in aCSF at the time of recording and bath applied for 10-20 mins before recordings commenced to allow for equilibration. All electrodes were calibrated to known DA standards in aCSF using a custom-made flow cell.

### Compounds

THP was purchased from Sigma-Aldrich (St. Louis, MO). Dihydro-β-erythroidine (DHβE) was obtained from Tocris (Minneapolis, MN). VU6021625 was generously provided by Dr. P. Jeffrey Conn (Vanderbilt University, TN).

### Statistical analysis

All data are presented as means with standard error. For all FSCV experiments, we used one slice per animal, so all samples are independent. Dose response experiments were analyzed with non-linear regression to determine EC_50_. EC_50_ was analyzed using a two-tailed Student’s *t*-test. Fluorescent *in situ* hybridization data was analyzed using one-way ANOVA with *post hoc* Dunnett’s multiple comparison test. Experiments assessing DA release were analyzed using 2-way repeated measures ANOVA (genotype x treatment) with *post hoc* Sidak’s multiple comparison test. Analyses were performed separately for *Tor1a*^*+/+*^ and *Tor1a*^*+/ΔE*^ in mAChR KO experiments. All analyses were performed using Graphpad Prism 9 (https://www.graphpad.com/). Statistical significance is defined as \**p* < 0.05, \*\**p* < 0.01, \*\*\**p* < 0.001, \*\*\*\**p* < 0.0001.

## RESULTS

### M4 mAChRs in striatal cholinergic cells mediate the effects of THP

We have previously demonstrated that the non-selective mAChR antagonist THP enhances DA release in both *Tor1a*^*+/+*^ and *Tor1a*^*+/ΔE*^ KI mice and rescues the deficiency in DA release in *Tor1a*^*+/ΔE*^ KI mice (Downs et al., 2019). However, the mAChR subtype(s) and cell types that mediate this effect are unknown. M4 mAChRs expressed on dSPNs, ChIs, and cortical glutamatergic terminals are known to regulate striatal DA release (Harrison et al., 1996; Threlfell et al., 2010; Threlfell and Cragg, 2011; Pancani et al., 2014). Therefore, to determine if THP depends on M4 mAChRs on any of these cell types, we assessed striatal DA release in response to challenge with THP in *Tor1a*^*+/+*^ and *Tor1a*^*+/ΔE*^ KI mice lacking M4 mAChRs on ChIs (ChI-M4^-/-^) or M4 mAChRs on dSPNs (dSPN-M4^-/-^). Deletion of M4 mAChRs from cortical glutamatergic inputs was not investigated because we have previously demonstrated that blocking ionotropic glutamate receptors does not affect the ability of THP to enhance DA release (Downs et al., 2019).

Previous studies have demonstrated that M4 mAChRs on dSPNs regulate striatal DA release by modulating endocannabinoid production (Foster et al., 2016). To test the hypothesis that THP enhances DA release by acting on dSPNs, we measured electrically evoked DA release in striatal slices from mice lacking M4 mAChRs in direct SPNs (dSPN-M4^-/-^) in the presence of THP (300 nM). Male and female *Tor1a*^*+/+*^ mice had a similar response to THP, so male and female mice were combined for all studies (main effect of sex, F_1,8_ = 0.039, p = 0.85, treatment x sex interaction effect, F_1,8_ = 0.00012, p = 0.99). THP did not alter DA clearance (tau), which is a surrogate for reuptake, in either *Tor1a*^*+/+*^ or *Tor1a*^*+/ΔE*^ mice (main effect of treatment, F_1,6_ = 0.046, p = 0.84). The dSPN-M4 KO had no effect on baseline DA release in either *Tor1a*^*+/+*^ mice (Sidak’s multiple comparison test, p = 0.78) or *Tor1a*^*+/ΔE*^ (Sidak’s multiple comparison test, p = 0.99). Deleting M4 mAChR expression from dSPNs had no effect on the THP-induced increase in DA release in either *Tor1a*^*+/+*^ mice (main effect of treatment, F_1,7_ = 120.3, p < 0.0001; treatment x genotype interaction effect, F_1,7_ = 0.09, p = 0.76) or *Tor1a*^*+/ΔE*^ mice (**Fig. 1*A***, main effect of treatment, F_1,7_ = 91.38, p < 0.0001; treatment x genotype interaction effect, F_1,7_ = 1.68, p = 0.24).

**Figure 1.**
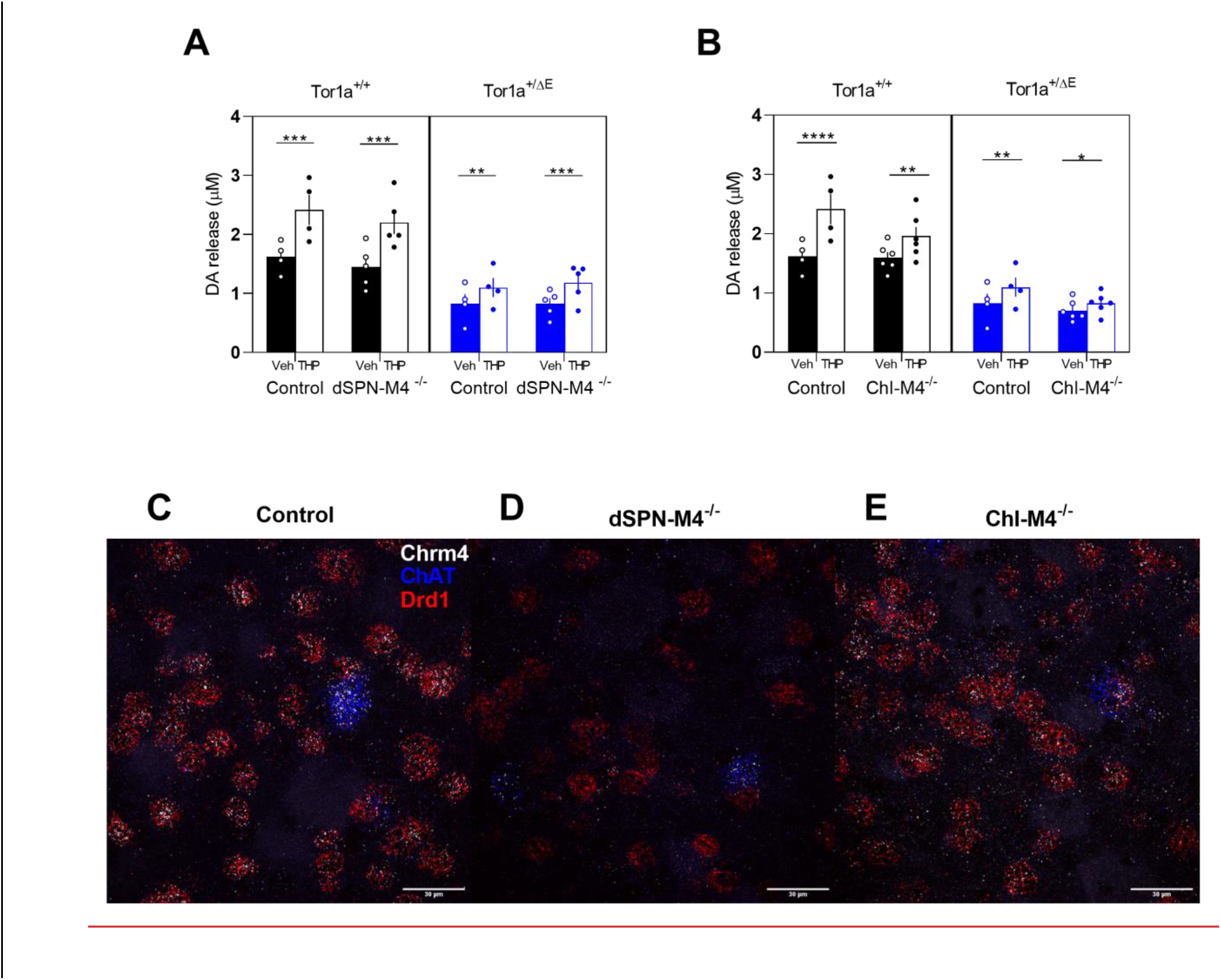
Effect of cell-type specific deletion of M4 mAChRs on striatal DA release in *Tor1a*^*+/+*^ and *Tor1a*^*+/ΔE*^ KI mice. ***A***. KO of M4 mAChR from direct SPNs did not significantly change the DA-release enhancing effects of 300nM THP in either genotype. ***B***. KO of M4 mAChR from ChIs reduced the effect of THP compared to control mice that were homozygous for the flox’d M4 mAChR allele but did not carry a cre transgene. Data are expressed as mean peak concentration of DA released (μM) ± SEM (n=4-6); *p<0.05, **p<0.01, ****p<0.0001. ***C, D, E***. Representative images of fluorescent *in situ* hybridization using probes to ChAT, Chrm4, and Drd1 mRNA in control ***C***., dSPN-M4^-/-^ ***D***., and ChI-M4^-/-^ ***E***. mice demonstrate cell-type specific deletion of M4 mAChR. Scale bar = 30 μm.

To verify that the M4 mAChR conditional knockout in dSPNs was both effective and cell-type specific, we used fluorescent *in situ* hybridization with probes to *Chrm4, Drd1* and *ChAT* to assess striatal mRNA expression. *Chrm4* encodes the M4 mAChR. *Drd1* encodes the D1 DA receptor which is a marker for dSPNs and *ChAT* encodes choline acetyltransferase, a marker for ChIs. In control mice, *Chrm4* mRNA was expressed in all *ChAT*-positive cells and almost all *Drd1*-positive cells, as expected. In dSPN-M4^-/-^ mice, there was a significant reduction in the percent of cells expressing both *Chrm4* and *Drd1* mRNA to ∼13% of control mice while there was a corresponding increase in the percent of cells that expressed only *Drd1* without *Chrm4* (**Fig 1***C* **vs *D &* Table 2**, one-way ANVOA, F_2,6_ = 554.7, p < 0.0001). There was no change in *Chrm4* mRNA expression in *ChAT*-positive striatal ChIs in dSPN-M4^-/-^ compared to control mice. These results validate the cell-type specific conditional knockout of M4 mAChRs in dSPNs but not ChIs.

**Table 2.**
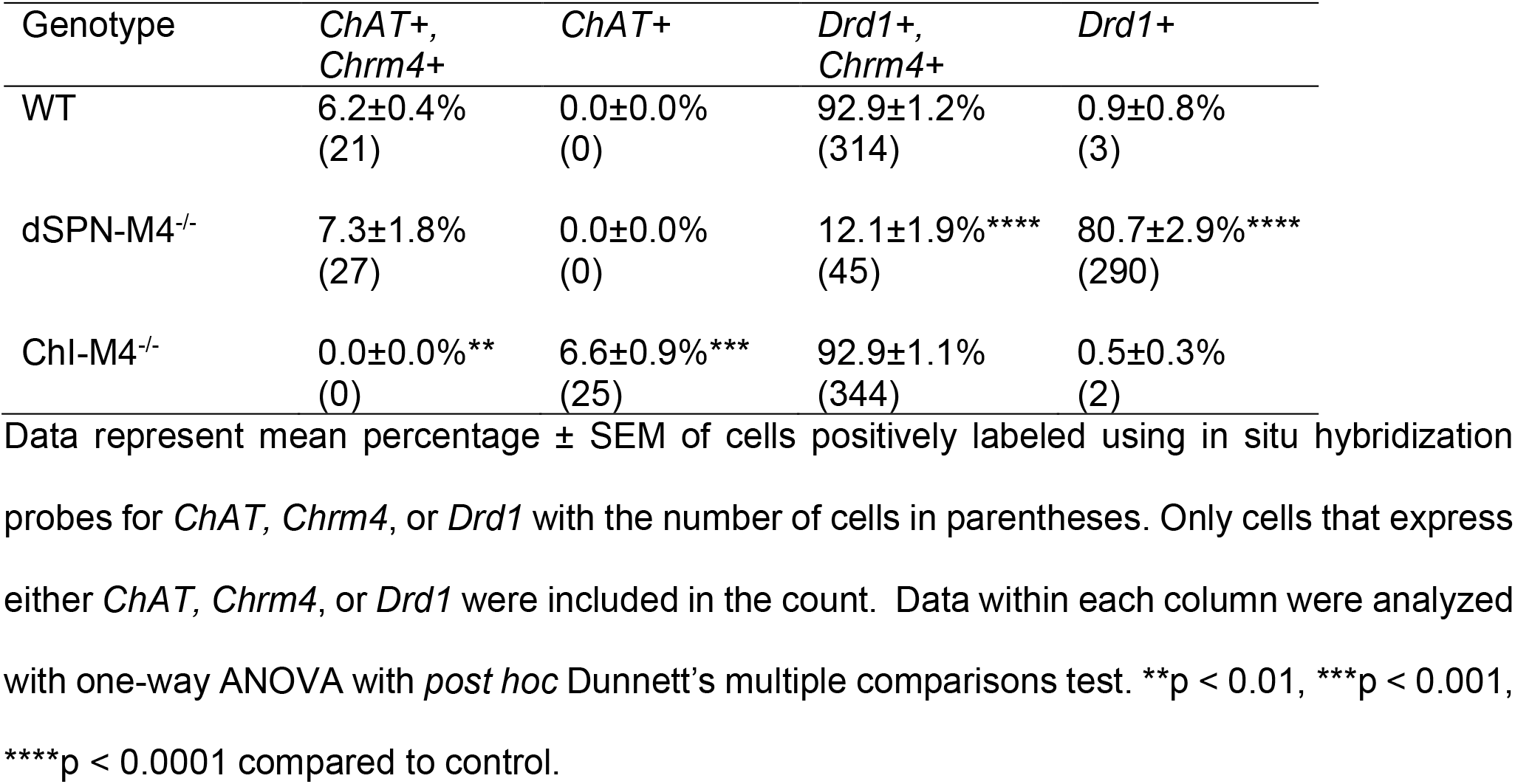
M4 mAChR KO mouse validation

M4 mAChRs are expressed on ChIs where they act as inhibitory autoreceptors. It is known that activation of these autoreceptors results in a reduction in DA release via a decrease in nicotinic acetylcholine receptor (nAChR) activation on DA terminals (Threlfell et al., 2010). To test the hypothesis that THP exerts its effect by blocking M4 mAChRs expressed on ChIs, we assessed DA release in striatal slices from mice lacking M4 mAChRs in ChIs (ChI-M4^-/-^). KO of M4 mAChRs from ChIs had no effect on baseline DA release in either *Tor1a*^*+/+*^ (Sidak’s multiple comparison test, p = 0.99) or *Tor1a*^*+/ΔE*^ mice (Sidak’s multiple comparison test, p = 0.32). While THP increased DA release in both *Tor1a*^*+/+*^ and *Tor1a*^*+/+*^; ChI-M4^-/-^ mice (**Fig 1*B***, main effect of treatment F_1,8_ = 64.22, p < 0.0001), the effect of THP was significantly attenuated in *Tor1a*^*+/+*^; ChI-M4^-/-^ mice to 65% of *Tor1a*^*+/+*^ mice (treatment x genotype interaction effect, F_1,8_ = 9.043, p = 0.017, Sidak’s multiple comparison test, p = 0.12). In *Tor1a*^*+/ΔE*^ mice, THP significantly increased DA release (main effect of treatment, F_1,8_ = 29.05, p = 0.0007; treatment x genotype interaction effect F_1,8_ = 1.60, p = 0.24). While THP increased DA release in *Tor1a*^*+/ΔE*^; ChI-M4^-/-^ mice (Sidak’s multiple comparisons test, p = 0.02), the effects of THP were attenuated from a 32% increase in DA release in control animals, to a 17% increase in *Tor1a*^*+/ΔE*^; ChI-M4^-/-^ mice. However, this decrease did not reach statistical significance (Sidak’s multiple comparisons test, p = 0.13). These results suggest that M4 mAChRs on ChIs mediate the effects of THP, in part, in both *Tor1a*^*+/+*^ and *Tor1a*^*+/ΔE*^ KI mice.

Fluorescent *in situ* hybridization with probes to *Chrm4, Drd1* and *ChAT* was used to verify that the M4 mAChR conditional knockout in ChIs was accomplished. In control mice, *Chrm4* mRNA was detected in all *ChAT*-positive cells. However, in ChI-M4^-/-^ mice, *Chrm4* mRNA expression was not detected in any *ChAT*-positive cells (**Fig 1*C*** **vs *E &* Table 2**, one-way ANVOA F_2,6_ = 18.68, p = 0.0002). There was no change in *Chrm4* mRNA expression in *Drd1-*positive cells in ChI-M4^-/-^ compared to control mice. Thus, the conditional *Chrm4* deletion targeted to ChIs was cell-type specific and highly effective.

### A selective M4 mAChR antagonist is sufficient to recapitulate the effects of THP

The cell-type selective KO experiments suggest that M4 mAChRs are necessary for the DA release-enhancing effects of THP in *Tor1a*^*+/ΔE*^ KI mice. However, because THP acts at all five mAChRs, it is not clear if selective blockade of M4 mAChRs would rescue DA release in *Tor1a*^*+/ΔE*^ KI mice similar to the effects of THP. Therefore, we assessed DA release in the presence of the subtype-selective M4 mAChR antagonist VU6021625, which was shown to be effective in ameliorating the dystonia exhibited by a mouse model of DOPA-response dystonia (Moehle et al., 2020). VU6021625 dose-dependently enhanced DA release in both *Tor1a*^*+/+*^ and *Tor1a*^*+/ΔE*^ KI mice (**Fig 2*A***). There was no significant difference in EC_50_ between genotypes (Student’s *t*-test, p = 0.87). VU6021625 did not alter DA clearance kinetics in either genotype (main effect of treatment, F_1,6_ = 0.002, p = 0.96. DA release was significantly increased in both *Tor1a*^*+/+*^ and *Tor1a*^*+/ΔE*^ KI mice at the maximal concentration of 300nM VU6021625 (**Fig 2*B***, main effect of treatment, F_1,8_ = 30.19, p = 0.0006). VU6021625 was similarly effective in both *Tor1a*^*+/+*^ and *Tor1a*^*+/ΔE*^ KI mice (genotype x treatment interaction effect, F_1,8_ = 1.37, p = 0.27). Importantly, VU6021625 normalized DA release in *Tor1a*^*+/ΔE*^ KI mice to *Tor1a*^*+/+*^ levels (Sidak’s multiple comparison’s test, p = 0.58).

**Figure 2.**
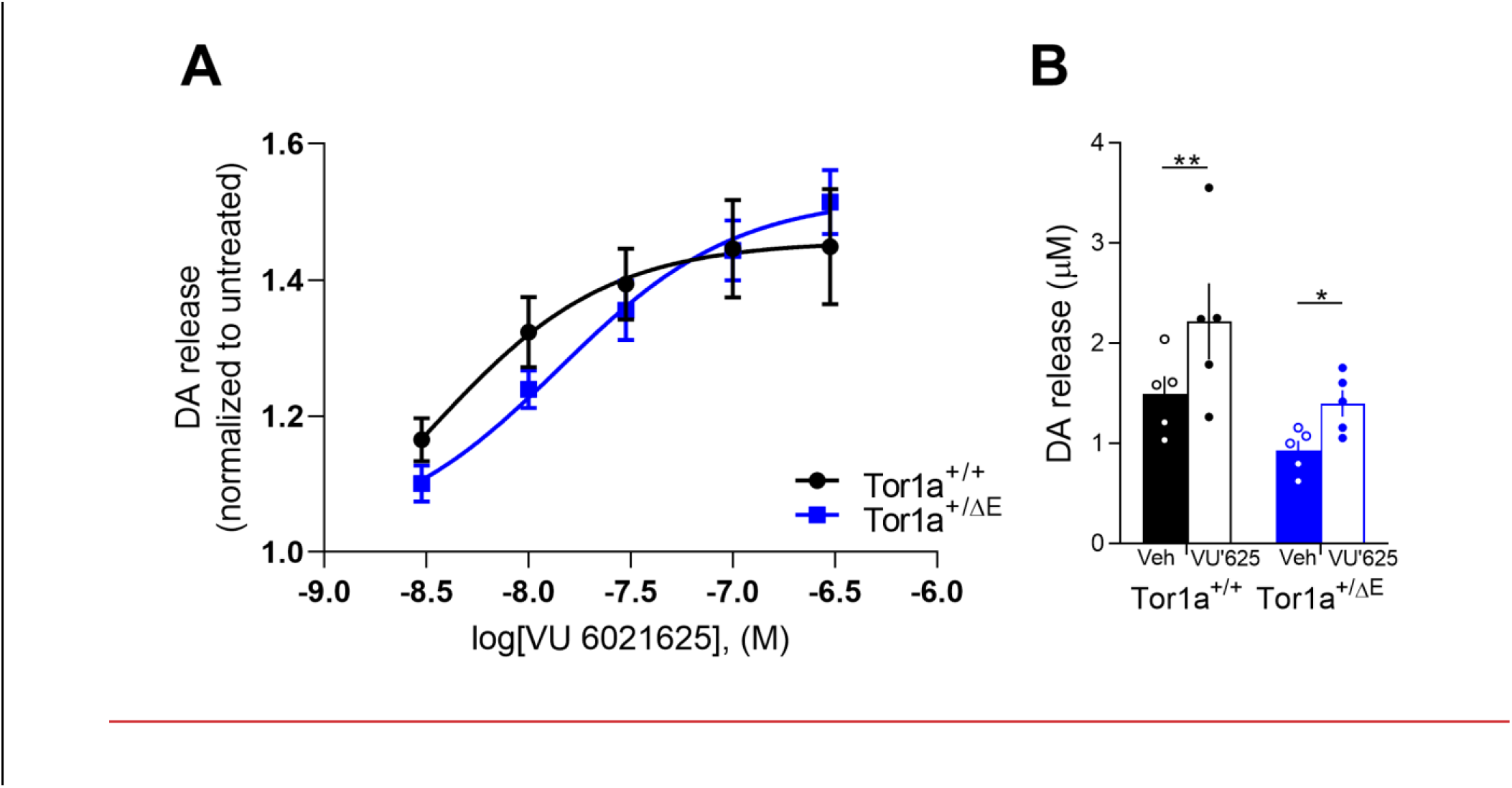
Effect of VU6021625 on striatal DA release in *Tor1a*^*+/+*^ and *Tor1a*^*+/ΔE*^ KI mice. ***A***. Dose response curve of the effect of the subtype selective M4 mAChR antagonist VU6021625 on DA release. VU6021625 dose-dependently enhanced DA release in both *Tor1a*^*+/+*^ and *Tor1a*^*+/ΔE*^ KI mice. Values are normalized to untreated for each genotype. ***B***. Effect of maximal concentration of VU6021625 on DA release. 300nM VU6021625 significantly enhanced striatal DA release in both *Tor1a*^*+/+*^ and *Tor1a*^*+/ΔE*^ KI mice. Values represent mean ± SEM (n=5); *p<0.05, **p<0.01.

### Blockade of M4 mAChRs in ChIs but not dSPNs blocks the effects of the M4 subtype selective antagonist VU6021625

To determine the specific cell type(s) that mediate the effects of the M4 mAChR subtype-selective antagonist, we assessed DA release in response to VU6021625 (300 nM) in dSPN-M4^- /-^ mice. Deleting M4 mAChR expression from dSPNs had no effect on the VU6021625-induced increase in DA release in *Tor1a*^*+/+*^ (main effect of treatment F_1,7_ = 121.0, p < 0.0001; treatment x genotype interaction effect, F_1,7_ = 0.66, p = 0.44). While VU6021625 enhanced DA release in both *Tor1a*^*+/ΔE*^ and *Tor1a*^*+/ΔE*^ dSPN-M4^-/-^ mice (**Fig 3*A***, main effect of treatment F_1,7_ = 134.0, p < 0.0001), VU6021625 appeared more effective at elevating DA release in *Tor1a*^*+/ΔE*^ dSPN-M4^-/-^ mice (treatment x genotype interaction effect, F_1,7_ = 24.74, p = 0.0016, Sidak’s multiple comparison test, p = 0.06). These results suggest that while M4 mAChR on dSPN are not required for the DA-enhancing effects of VU6021625 in *Tor1a*^*+/+*^ mice, M4 mAChR on dSPNs may inhibit DA release in *Tor1a*^*+/ΔE*^ KI animals.

**Figure 3.**
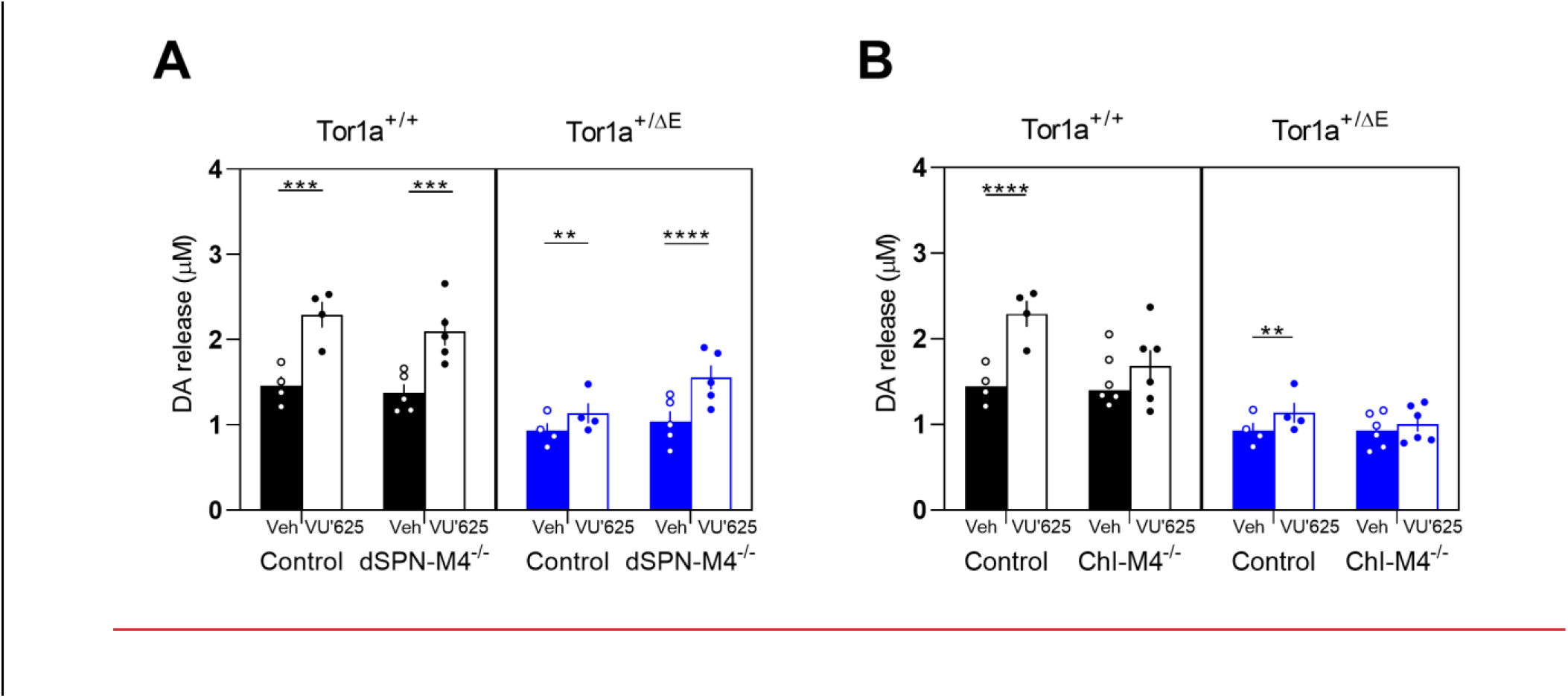
Effect of M4 mAChR KO in dSPNs or ChIs on the VU6021625-induced increase in DA release. ***A***. KO of M4 mAChR from dSPNs did not significantly affect the DA release-enhancing effects of 300nM VU6021625 compared to control mice that were homozygous for the flox’d M4 mAChR allele but did not carry a cre transgene. ***B***. KO of M4 mAChR from ChIs reduced the DA release enhancing effects of VU6021625 in both *Tor1a*^*+/+*^ and *Tor1a*^*+/ΔE*^ mice. Data are expressed as concentration of DA released (μM) ± SEM (n=4-6); **p<0.01, ****p<0.0001.

Next, we determined the role of M4 mAChR expressed on ChIs in mediating the increase in DA release in response to VU6021625 treatment. KO of M4 mAChR from ChIs eliminated the DA enhancing effect of VU6021625 in *Tor1a*^*+/+*^ mice (**Fig 3*B***, treatment x genotype interaction effect, F_1,8_ = 31.29, p = 0.0005, Sidak’s multiple comparison test, p = 0.16). KO of M4 mAChR from cholinergic neurons also eliminated the DA enhancing effect of VU6021625 in *Tor1a*^*+/ΔE*^ KI mice (treatment x genotype interaction effect, F_1,8_ = 8.85, p = 0.017, Sidak’s multiple comparison test, p = 0.053). These data suggest that the M4 subtype selective mAChR antagonist enhances DA release by blocking M4 mAChR on ChIs in both *Tor1a*^*+/+*^ and *Tor1a*^*+/ΔE*^ KI mice.

### M1 mAChRs on dSPNs and iSPNs are not required for the effects of THP in Tor1a^+/+^ or Tor1a^+/ΔE^ KI mice

M1 mAChRs are expressed on both direct and indirect SPNs (Harrison et al., 1996; Hernandez-Flores et al., 2015). Previous studies have demonstrated that M1 mAChR regulates extracellular DA in mice (Gerber et al., 2001a). Although the mechanism mediating this effect is largely unknown, it is known that SPNs provide GABAergic input to ChIs (Tepper et al., 2008). It is possible that THP mediates DA release indirectly by blocking excitatory M1 mAChRs on SPNs to disinhibit ChI activity thereby increasing ACh release and nAChR activity on DA terminals. To determine if M1 mAChRs on either direct or indirect SPNs are required for the DA release-enhancing effects of THP, we assessed striatal DA release in response to challenge with THP in *Tor1a*^*+/+*^ and *Tor1a*^*+/ΔE*^ KI mice lacking M1 mAChR on dSPNs (dSPN-M1^-/-^) or iSPNs (iSPN-M1^-/-^).

In *Tor1a*^*+/+*^ mice lacking M1 mAChR in dSPNs, THP significantly increased DA release in both *Tor1a*^*+/+*^ and *Tor1a*^*+/+*^ dSPN-M1^-/-^ mice (**Fig 4A** treatment effect, F_1,8_ = 63.92, p < 0.0001, Sidak multiple comparisons test, p = 0.0004). Although there was a trend towards reduced efficacy in *Tor1a*^*+/+*^ dSPN-M1^-/-^ mice (treatment x genotype interaction effect, F_1,8_ = 4.63, p = 0.064), KO of M1 mAChR from dSPNs did not affect the response to THP in *Tor1a*^*+/ΔE*^ KI mice (treatment x genotype interaction effect F_1,8_ = 0.22, p = 0.65). There were no differences observed in baseline DA release in either *Tor1a*^*+/+*^ (Sidak’s multiple comparison test, p = 0.98) or *Tor1a*^*+/ΔE*^ (Sidak’s multiple comparison test, p = 0.57) due to dSPN-M1 KO. These results demonstrate that M1 mAChRs on dSPNs do not significantly contribute to the effects of THP on DA release in either *Tor1a*^*+/+*^ or *Tor1a*^*+/ΔE*^ KI mice.

**Figure 4.**
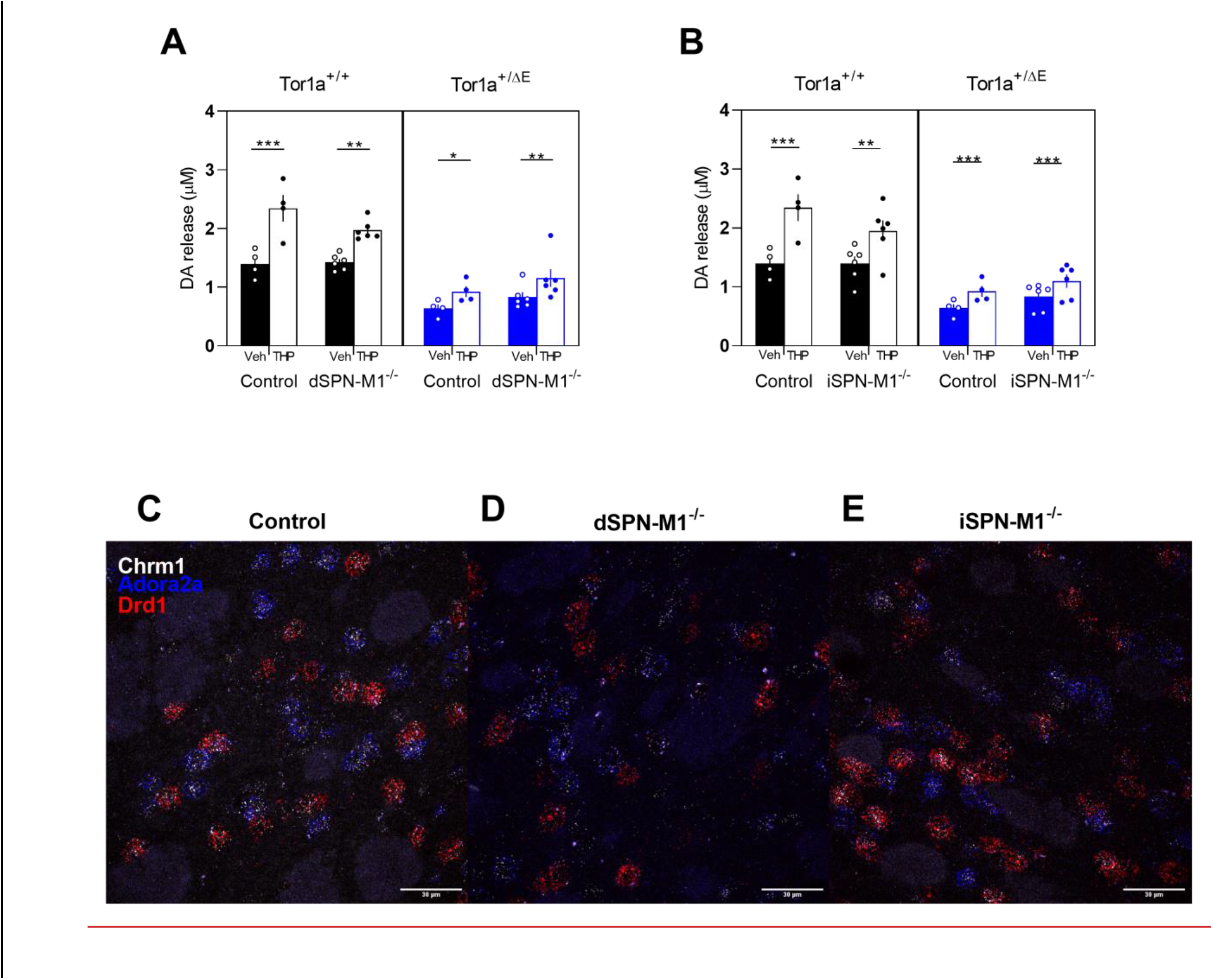
Effect of cell-type specific deletion of M1 mAChRs on striatal DA release in *Tor1a*^*+/+*^ and *Tor1a*^*+/ΔE*^ KI mice. ***A***. KO of M1 mAChR from direct SPNs did not alter the effects of THP in *Tor1a*^*+/+*^ or *Tor1a*^*+/ΔE*^ KI mice compared to control mice that were homozygous for the flox’d M1 mAChR allele but did not carry a cre transgene. ***B***. KO of M1 mAChR from indirect SPNs did not alter the effects of THP in *Tor1a*^*+/+*^ or *Tor1a*^*+/ΔE*^ KI mice compared to control mice that were homozygous for the flox’d M1 mAChR allele but did not carry a cre transgene. Data are expressed as concentration of DA released (μM) ± SEM. (n=4-6); *p<0.05, **p<0.01, ***p<0.001, ****p<0.0001. Representative images of control ***C***., dSPN-M4^-/-^ ***D***., and iSPN-M4^-/-^ ***E***. mice labeled for ChAT, Chrm4, and Drd1 mRNA. Scale bar = 30 μm.

In *Tor1a*^*+/+*^ mice lacking M1 mAChR in iSPNs, THP significantly increased DA release (**Fig 4*B***, main effect of treatment, F_1,8_ = 76.78, p < 0.0001). However, there was a trend towards reduced efficacy in *Tor1a*^*+/+*^ iSPN-M1^-/-^ mice (treatment x genotype interaction effect, F_1,8_ = 5.24, p = 0.051). The effect of THP in *Tor1a*^*+/ΔE*^ iSPN-M1^-/-^ mice did not differ from *Tor1a*^*+/ΔE*^ mice (treatment x genotype interaction effect F_1,8_ = 0.147, p = 0.71). Deletion of M1 mAChRs from iSPNs had no effect on baseline release in either *Tor1a*^*+/+*^ (Sidak’s multiple comparison test, p < 0.99) or *Tor1a*^*+/ΔE*^ (Sidak’s multiple comparison test, p < 0.43). These results demonstrate that M1 mAChRs on iSPNs do not significantly contribute to the effects of THP on DA release in either *Tor1a*^*+/+*^ or *Tor1a*^*+/ΔE*^ KI mice.

To confirm that the SPN cell-type specific KO of M1 mAChRs were effective, we used fluorescent *in situ* hybridization with probes specific to *Chrm1, Adora2a*, and *Drd1. Chrm1* encodes the M1 mAChR receptor. *Adora2a* encodes the A2A adenosine receptor and is a specific marker for iSPNs. *Drd1* is a specific marker for dSPNs. In control mice, *Chrm1* was expressed in almost all *Adora2a*-positive and *Drd1*-positive cells. However, in dSPN-M1^-/-^ mice, the percentage of cells expressing both *Chrm1* and *Drd1* mRNA was significantly reduced to ∼2% of control levels (**Fig 4*C*** **vs *D &* Table 3**, one-way ANVOA F_2,6_ = 90.44, p < 0.0001) while the percentage of cells expressing both *Chrm1* and *Adora2a* was unchanged. In iSPN-M1^-/-^mice, the percentage of cells expressing both *Chrm1* and *Adora2a* mRNA was significantly reduced to ∼5% of control levels (**Fig 4*C*** vs *E &* Table 3, one-way ANOVA F_2,6_ = 57.37, p < 0.0001), while the percentage of cells expressing both *Chrm1* and *Drd1* mRNA was unchanged. These results demonstrate that the M1 mAChR knockouts were specific and effective.

**Table 3.** M1 mAChR KO mouse validation

| Genotype | *Adora2a*+,<br>*Chrm1*+ | *Adora2*+ | *Drd1*+,<br>*Chrm1*+ | *Drd1*+ |
| --- | --- | --- | --- | --- |
| Control | 47.4±4.0%<br>(213) | 3.2±0.6%<br>(16) | 49.1±3.1%<br>(226) | 0.2±0.2%<br>(1) |
| dSPN-M1<sup>-/-</sup> | 46.5±3.4%<br>(220) | 4.0±0.3%<br>(20) | 2.3±0.6%****<br>(11) | 46.9±3.3%****<br>(223) |
| iSPN-M1<sup>-/-</sup> | 4.4±1.0%***<br>(23) | 42.3±3.2%****<br>(224) | 52.8±3.7%<br>(279) | 0.4±0.4%<br>(2) |

Data represent mean percentage ± SEM of *Chrm1*, *Adora2a*, and *Drd1* positive cells (number of cells). Only cells that express either *Chrm1*, *Adora2a*, or *Drd1* were included in the count. Data within each column were analyzed with one-way ANOVA with *post hoc* Dunnett's multiple comparisons test. \*\*\*p < 0.001, \*\*\*\*p < 0.0001 compared to Control.

## DISCUSSION

THP is the only oral medication to be proven effective in a double-blind placebo-controlled trial for dystonia (Fahn, 1983; Burke et al., 1986). Unfortunately, most patients cannot use THP due to intolerable side-effects at the high doses necessary for efficacy in dystonia (Schwarz and Bressman, 2009; Thenganatt and Jankovic, 2014). These side effects include cognitive impairment, constipation, dry mouth, urinary retention, and others (Burke et al., 1986; Jabbari et al., 1989; Guthrie et al., 2000; Lumsden et al., 2016). Here we used THP as a platform to further understand the mechanism of action of mAChR antagonists and identify specific mAChR subtypes that may serve as targets for more specific therapeutics with fewer side effects. We demonstrated that M4 mAChR located on striatal ChIs mediate, in part, the THP-induced increase in striatal DA release. Furthermore, we found that M1 mAChRs likely do not play a role in the effects of THP.

Our studies demonstrate that M1 mAChRs on neither dSPNs nor iSPNs contribute to the effects of THP in *Tor1a*^*+/+*^ or *Tor1a*^*+/ΔE*^ KI mice. This is in contrast to a previous study that found elevated extracellular DA in the striatum in global M1 mAChR KO mice, albeit in an *in vivo* setting using microdialysis (Gerber et al., 2001b). This may reflect the fact that long loop connections are severed in our *ex vivo* slice preparation. Further, any local-circuit effects of M1 mAChR activation are likely to be indirect perhaps via SPN axon collateral synapses onto ChIs (Tepper et al., 2008). These connections might not be functional in our slice preparation or may be occluded by the local electrical stimulation used in our study.

Our results demonstrate that M4 mACHRs, in part, mediate the THP-induced rescue of DA release in *Tor1a*^*+/ΔE*^ KI mice. KO of M4 mAChR from ChIs, but not dSPNs, diminished the increase in DA release caused by THP in both *Tor1a*^*+/+*^ and *Tor1a*^*+/ΔE*^ KI mice. The residual effect of THP on DA release may be a result of THP acting on M2 mAChRs also located on striatal cholinergic interneurons, which are presumed to have an identical function to M4 mAChRs located on these cells (Threlfell et al., 2010). This is supported by the fact that KO of M4 from ChIs abolishes the effects of the M4 mAChR selective antagonist VU6021625. Surprisingly, KO of M4 mAChR from ChIs by itself did not normalize DA release as illustrated by the comparable baselines between control and KO mice. This may be due to homeostatic changes during development. In this context, the results from the antagonist experiments in intact adult mice are particularly important, because gene deletion experiments can be confounded by compensatory developmental changes in response to the knockout. Further, the fact that KO of M4 MAChR from ChI abolished the effects of VU6021625 in both *Tor1a*^*+/+*^ and *Tor1a*^*+/ΔE*^ KI mice demonstrates the specificity of this compound. However, the response to the M4 mAChR antagonist in adult mice mirrored the response in the KO mice. Thus, the consistency of our results across both the M4 mAChR antagonist challenge in genetically unaltered adult mice and the conditional knockout experiments suggests that developmental confounds do not likely account for the results.

Although *Tor1a*^*+/ΔE*^ KI mice do not exhibit a dystonic phenotype (Goodchild et al., 2005) these mice do exhibit a reduction in DA neurotransmission, a phenotype observed across many different forms of dystonia. Thus, the ability of THP and the subtype-selective mAChR antagonists to normalize striatal DA release in *Tor1a*^*+/ΔE*^ KI mice represents a plausible mechanism of action for these compounds. Indeed, we have previously demonstrated that THP and the M4 mAChR subtype-selective antagonist VU6021625 ameliorates the dystonic movements exhibited by a knockin mouse model of DOPA-responsive dystonia, another model in which DA neurotransmission is significantly reduced (Rose et al., 2015; Moehle et al., 2020). Abnormalities in striatal DA neurotransmission have been implicated in diverse forms of dystonia including, adult-onset idiopathic dystonias, secondary dystonia due to Parkinson’s disease or typical antipsychotic treatment, and inherited dystonia (Ichinose and Nagatsu, 1999; Tolosa and Compta, 2006; Carbon et al., 2009; Berman et al., 2013; Rilstone et al., 2013; Ng et al., 2014; Mehta et al., 2015). Thus, abnormal striatal DA neurotransmission is a convergent pathology across dystonias with diverse etiologies. While DA receptor agonists are generally not considered to be effective in dystonia, a recent reappraisal of these therapeutics suggests that both direct and indirect DA agonists are effective in subsets of dystonia patients and mouse models of dystonia (Fan et al., 2018). Further, unlike direct DA receptor agonists, which act at DA receptors in a temporally indiscriminate manner, THP and other mAChR antagonists provide the advantage of enhancing DA release while maintaining normal patterns of DA release encoded by DA neuron firing. However, mAChR antagonists may act through mechanisms in addition to DA neurotransmission to diminish dystonic symptoms. Previous studies have demonstrated that THP normalizes abnormal corticostriatal plasticity, including diminished long-term depression and loss of synaptic depotentiation in SPNs, observed in *Tor1a*^*+/ΔE*^ KI mice via M1 mAChRs (Maltese et al., 2014; Martella et al., 2014). Thus, it is possible that a multiprong approach may be most effective for the amelioration of dystonia.

The role of ChIs in the mechanism of action of THP is consistent with the considerable evidence implicating ChI dysfunction in *TOR1A* dystonia. Normally, D2 DA receptor activation inhibits ChI firing rates, causing a reduction in extracellular ACh. In contrast, in *Tor1a(ΔE)* rodent models, the D2 DA receptor agonist quinpirole induces an increase in the ChI firing rate (Pisani et al., 2006; Martella et al., 2009; Grundmann et al., 2012; Sciamanna et al., 2012). Both THP and a more selective M2/M4 mAChR antagonist restore D2 DA receptor-mediated ChI responses in *ex vivo* brain slices of *Tor1a*^*+/ΔE*^ KI mice (Scarduzio et al., 2017). Thus, THP and other more selective mAChR antagonists normalize both DA release and striatal ChI responses in *Tor1a*^*+/ΔE*^ KI mice suggesting a central role for ChI in the development of novel therapeutics. However, it is not yet known if a selective M4 mAChR antagonist alone would similarly normalize ChI responses.

Taken together, our data demonstrate that THP normalizes DA release in *Tor1a*^*+/ΔE*^ KI mice by blocking M4 mAChR on ChIs. This suggests that M4 subtype-selective mAChR antagonists, alone or in combination with other compounds, may be promising therapeutics for *TOR1A* dystonia. Further, because many of the dose-limiting peripheral side effects associated with THP, including constipation, dry mouth, and urinary retention, are mediated by M2 and M3 mAChR (Bymaster et al., 2003), M4 subtype-selective mAChR antagonists may improve dystonia while avoiding many of the side effects of the non-selective mAChR antagonists that are currently available to patients.

## Abbreviations

ACh: acetylcholine
aCSF: artificial cerebrospinal fluid
ChI: cholinergic interneuron
DA: dopamine
dSPN: direct striatal projection neuron
FSCV: fast scan cyclic voltammetry
iSPN: indirect striatal projection neuron
KI: knockin
KO: knock out
mAChR: muscarinic acetylcholine receptor
nACHR: nicotinic acetylcholine receptor
THP: trihexyphenidyl

